# Isolation and characterization of fucosylated extracellular vesicles based on a novel high-throughput GlyExo-Capture technique

**DOI:** 10.1101/2021.12.09.471505

**Authors:** Boan Li, Kun Hao, Cuidie Ma, Zhengtai Li, Hongjiang Li, Wenqian Du, Lijuan Sun, Tianye Jia, Aixia Liu, Yanzhao Li, Lida Xu, Qi Gao, Ruifu Yang, Changqing Lin

**Author notes:** Correspondence (L.X.), (Q.G.), (R.Y.), (C.L.). These authors contributed equally to the work (Boan Li, Kun Hao).

## Abstract

Owing to their diagnostic and therapeutic potential, extracellular vesicles (EVs) derived from tumour cells have recently garnered great interest. The presence of different glycosylation sites at the EV surface supports the need for efficient glycosylated EV isolation. Here, we developed a GlyExo-Capture technique for robustly capturing fucosylated EVs from sera and cell supernatants. Lens culinaris lectin (LCA)-immobilized magnetic complexes were found to capture approximately 60% of the total EVs from HepG2 cells. The capture efficiency was reduced to less than 40% in nontumorigenic MIHA cells. Notably, the cellular uptake pattern of highly fucosylated EVs was markedly different from that of EVs with low fucosylation. The unearthing of enriched fucosylated EV miRNA cargos by next-generation deep sequencing (NGS) revealed 75 differentially expressed miRNAs (DEMs) in hepatocellular carcinoma (HCC). Among them, a 4-miRNA panel was chosen and yielded an area under the ROC curve (AUC) of 0.86 and 0.84 for the detection of HCC from non-HCC controls in testing samples and independent validation samples, respectively. The 4-miRNA signature was independent of alpha-fetoprotein (AFP), and a combined model with AFP yielded an increased AUC of 0.92. In conclusion, we developed a high-throughput method for capturing fucosylated EVs efficiently and shed light on the use of fucosylated EVs as potential sources of miRNAs for cancer biomarker detection.

## Introduction

Extracellular vesicles (EVs) are nanosized particles with lipid bilayer membranes that carry proteins, DNA, RNA, and other metabolites (1). EVs are capable of encapsulating and transporting several molecules for targeted delivery and play various roles in intercellular communication, normal physiological processes, and tumour development (2-4). EVs are widely present in human bodily fluids and can be secreted by various types of cells (5). Compared with nontumor cells, tumour cells have different amounts and compositions of EVs (6). Tumour cells always have a ‘ sweeter’ surface, and tumour-associated glycan alterations have already been identified as enriched in cancer EVs (7,8). Recent studies have shown that small EVs from non-small-cell lung carcinoma (NSCLC) cells are dominated by typical lung-type N-glycans with NSCLC-associated core fucosylation (9). These glycan-coated EVs can be described as ‘ sugar-coated bullets’ fired by tumour cells (10). The presence of different glycosylation profiles under normal and cancer conditions highlights the importance of studying glycosylated EVs and their specific miRNA cargos in cancer diagnosis and therapy (7,11-13).

Traditional EV isolation methods, which include ultracentrifugation, microfiltration, and size exclusion chromatography, are based on EV size and buoyant density (14). Recent immune affinity or microfluidic technologies are based on the exposed components on the EV surface. Owing to their abundance on the EV surface, glycans constitute a valuable source of targets to capture for isolating EVs from a heterogeneous sample. For example, lectins including peanut agglutinin (PNA), *Artocarpus integrifolia* (AIA or Jacalin) lectin, and *Maackia amurensis* lectin I (MAL-I) are used to isolate EVs of different sizes from human urine samples based on the high affinity of these lectins for Gal or GalNAc residues with a (2,3)-linked sialic acid (7). A similar approach was applied to isolate EVs from urine samples by *Solanum tuberosum* lectin (STL) based on its interaction with N-acetylglucosamine and lactosamine residues (7). Nanoparticle-based time-resolved fluorescence immunoassays (NP-TRFIAs) can capture EVs from urine samples and cell supernatants based on the tetraspanins and glycan antigens present on the EV surface (15). Apart from affinity-based isolation methods, commercially available precipitation kits (i.e., ExoGAG) can be used to precipitate EVs out of the solution due to the presence of negatively charged GAG at the EV surface (16). Despite technological advances in this field, less laborious isolation methods are still technically challenging.

In this study, we developed a GlyExo-Capture technique aiming to capture EVs by using the fucose-specific lens culinaris lectin (LCA) for affinity isolation. This GlyExo-Capture technique enriches fucosylated EVs from serum samples and cell supernatants over a short period and with high throughput. The characterization of enriched fucosylated EVs and the diagnostic value of fucosylated EV-specific miRNAs are also demonstrated, with the potential to be translated into the clinical setting.

## Material and Methods

### Patient enrolment and study cohorts

The study was approved by the Ethics Committee of the Fifth Medical Center of Chinese PLA General Hospital (2016-004-D). The patients included in the study were >18 years of age. Exclusion criteria were pregnancy, immunosuppression, leukopenia, presence of haematological malignancies, or initiation of palliative care. All patients with hepatocellular carcinoma (HCC) were treatment-naïve at the time of blood sampling. Three independent cohorts were included. The demographic and clinicopathologic characteristics of the patients in the three cohorts are shown in Supplementary Table S1.

1. Disclosure of miRNA profiles derived from fucosylated EVs by next-generation deep sequencing (NGS) was conducted for a cohort comprising 61 patients with HCC, 24 patients with hepatitis and 30 healthy subjects. The HCCs of more than 45% of the patients (47.5%, 29/61) were in the early stage. Healthy subjects and patients with hepatitis were included as the non-HCC group. This cohort, referred to as the original sample, was randomly divided into a training sample and a testing sample at a ratio of approximately 7:3. The training sample was used to construct prediction models based on the 4-miRNA signature in fucosylated EVs, while the testing sample was used to measure the prediction efficiencies of the models.
2. An independent cohort comprising 16 patients with HCC (50% of HCCs were in the early stage), 16 patients with hepatitis and 16 healthy subjects was included in this study to confirm the NGS data via real-time quantitative (PCR RT–qPCR).
3. An independent cohort comprising 30 patients with HCC, 52 patients with hepatitis and 23 healthy subjects was included as a validation sample to evaluate the 4-miRNA-based diagnostic potential. In this cohort, the HCCs of more than 45% of the with (46.7%, 14/30) were in the early stage.

### Sample collection and EV purification

Blood was drawn via venipuncture using a 21-gauge needle. To prevent haemolysis, aspiration was performed slowly and evenly for both procedures. Blood was collected in 5 mL vacuum tubes a containing separator gel (Improve Medical Instruments Co., Ltd, Guangzhou, China) and centrifuged at 1800 × g for 10 min at room temperature (RT) within 1 hour of sampling. The resulting sera were centrifuged again at 3000 × g for 10 min at 4 °C to remove any cellular debris. Then, the sera were aliquoted and stored at −80 °C. The EVs from sera samples were isolated by GlyExo-Capture according to a standard protocol.

### Reagents and resources

The key reagents and resources used in this research are listed in Supplementary Table S2.

### Cell culture and EV purification

HepG2 (CRL-10741), HCT-116 (CCL-247), and HeLa (CCL-2) cell lines were obtained from the American Type Culture Collection (ATCC, Rockville, USA). The MIHA (CE18544) cell line was obtained from Beijing Crisprbio Biotechnology Co., Ltd. All the cell lines were tested for mycoplasma contamination and were expanded in DMEM supplemented with 10% (v/v) foetal bovine serum (FBS; GIBCO, Life Technologies, Carlsbad, USA), 0.1 mg/mL streptomycin, 0.25 μg/mL amphotericin B and 100 U/mL penicillin (Sangon Biotech Co., Ltd., Shanghai, China) at 37 °C with 5% CO_2_. For EV production, the HepG2 cells were maintained in EV-depleted media, and the conditioned media were collected after 72 hours and then centrifuged at 300 × g for 10 min to remove the cells.

The cell supernatant was weak centrifuged at 3000 × g for 10 min to remove cellular debris. The resulting supernatant was then divided into two portions and transferred to new tubes. One portion was ultracentrifuged at 100,000 × g using a XPN-100 type 100 TI rotor (CP100NX; Hitachi, Brea, USA) for 90 min, and the resulting EV pellet was resuspended in 250 μL of cold PBS; these were denoted as UC-EVs. The other portion was isolated by GlyExo-Capture according to a standard protocol. In brief, it was incubated in the presence of LCA-immobilized beads, and the bound fraction was denoted as highly fucosylated EVs (HF-EVs). The unbound fraction was subjected to ultracentrifugation at 100,000 × g for 90 min, and the resulting EV pellet was denoted as constituting low fucosylated EVs (LF-EVs). To test the specificity of the GlyExo-Capture kit, 20 μg of EVs was treated with 2 μL of PNGase F (882 U/μL, Yeasen Biotechnology Co. Ltd., Shanghai, China) for 12 hours before a cellular uptake assay was performed. The protein content of EVs was quantified using a Qubit Protein Assay (Invitrogen, Pittsburgh, USA) and a Qubit 3.0 fluorometer (Invitrogen, Pittsburgh, USA).

### Transmission electron microscopy (TEM)

EVs isolated by ultracentrifugation GlyExo-Capture were fixed with 2% glutaraldehyde for 1 hour at RT and then stained with 0.5% uranyl acetate (pH 4.5). TEM was performed on an FEI Tecnai Spirit (FEI, Eindhoven, NL) at 120 kV. Electron micrographs were captured with a Gatan UltraScan 1000 charge-coupled device (CCD) camera (Pleasanton, USA).

### Nanoparticle tracking analysis (NTA)

The EVs isolated by ultracentrifugation or GlyExo-Capture were analysed for particle concentration and size distribution using a Malvern NanoSight NS300 instrument (Malvern Instruments, Ltd., Malvern, UK) in accordance with the manufacturer’ s instructions. Briefly, three independent replicates of diluted EV preparations in PBS were injected at a constant rate into the tracking chamber using the provided syringe pump. The specimens were tracked at RT for 60 s. The shutter speed and gain were adjusted and manually maintained at the optimized settings for the all samples. The data were captured and analysed with NTA Build 127 software version 2.3.

### Western blot analysis

EVs isolated by ultracentrifugation or GlyExo-Capture were lysed, and the protein concentration was analysed using a Qubit Protein Assay (Invitrogen, Pittsburgh, USA) and a Qubit 3.0 fluorometer (Invitrogen, Pittsburgh, USA). The input for EVs was normalized to 25 μg of total protein. The proteins were separated by SDS–PAGE and transferred to a 0.45 μm nitrocellulose membrane (GE Healthcare Life Sciences, Marlborough, USA); the membranes were blocked with 1% nonfat milk powder and incubated together with primary antibodies. The secondary antibody was rabbit anti-mouse IgG-HRP, and the protein bands were detected using a BeyoECL Plus Detection System (Beyotime Biotechnology, Shanghai, China). The primary antibodies used were anti-CD81 (M38) (1:400 dilution, Abcam, Cambridge, USA), anti-Alix (3A9) (1:400 dilution, Abcam, Cambridge, USA), anti-TSG101 (4A10) (1:400 dilution, Abcam, USA) and anti-calnexin (EPR3633 (2)) (1:2000 dilution, Abcam, Cambridge, USA).

### ExoView analysis

EVs isolated by ultracentrifugation or GlyExo-Capture were added to chips printed with antibodies including anti-human CD81 (JS-81), anti-human CD63 (H5C6), anti-human CD9 (HI9a), and anti-mouse IgG1 (MOPC-21) on an ExoView R100 instrument (NanoView Biosciences, Brighton, USA). All the chips were imaged by an ExoView scanner (NanoView Biosciences, Brighton, USA), and the data were analysed using NanoViewer 2.8.10 software (NanoView Biosciences, Brighton, USA).

### Cellular uptake assay via flow cytometry

The cellular uptake of HF-EVs, HF-EVs treated with PNGase F, LF-EVs, and LF-EVs treated with PNGase F was evaluated using flow cytometry. Briefly, the cells were exposed to DiD-labelled EVs at select doses and then incubated at 37 °C for the indicated time points. After extensive washing, flow cytometry was performed with a BD FACSCanto II instrument (BD Biosciences, San Jose, USA), and 45 s acquisitions under a medium flow rate were used for data capture. Data analysis was performed using FlowJo software with gating of cells in the PE channel, and no impact was found for equivalent DiD background samples. A control sample of the DiD dye background was prepared according to the same procedures as that indicated above but excluding cells.

### RNA isolation and analysis

Total RNA from EVs was extracted using a miRNeasy® Mini Kit (Cat. 217,004, Qiagen, Hilden, Germany) according to a standard protocol and was assessed using the High Sensitivity RNA Cartridge of a Qsep100 fully automatic nucleic acid analysis system (Bioptic, Inc., LA, USA).

### Next-generation sequencing and bioinformatics analysis

The EV-RNA was compiled into cDNA libraries using a NEBNext Small RNA Library Prep Set for Illumina (Multiplex Compatible) (Cat No.: E7330, NEB, Ipswich, USA) following the manufacturer’ s instructions. Library size was selected using the E-Gel SizeSelect II gel of an E-Gel Power Snap electrophoresis system (Thermo Fisher Scientific, Inc., Waltham, USA). The quality and concentration of the cDNA libraries were checked, and finally, groups of 24-26 samples were pooled at the same concentration before sequencing in an Illumina NextSeq 550 Sequencing System (75 nt, single read). The sequences were then assessed for quality, and primer-adapter sequences were trimmed by Cutadapt software, followed by their alignment to the human reference genome (HG38) by Bowtie. The aligned data were annotated by HT-seq with the gff3 file from miRBase version 23. The number of reads for each miRNA was normalized to transcripts per million (TPM) across all samples. The data have been uploaded to the Sequence Read Archive (SRA) database under BioProject number PRJNA785786.

The miRNA count data were normalized, and differential analysis was conducted using the DESeq2 package pf R 4.1.0. All the tests were two-tailed, and the false discovery rate (FDR) was used for multiple comparisons. Fold changes (FCs) larger than 2 and adjusted P values less than 0.05 (|log2(FC)|>1, p<0.05) were used to select differentially expressed miRNAs (DEMs) from a volcano plot. Heatmap (version 1.0.12) was used for clustering with the Euclidean distance method. Linear discriminant analysis (LDA) of the TPM expression matrix was conducted using the MASS package (version 7.3) to visualize the separation of HCC and non-HCC groups. Target genes of DEMs were predicted using miRPath on the DIANA website (http://snf-515788.vm.okeanos.grnet.gr/).

### RT–qPCR

cDNA was synthesized using a miRcute plus miRNA cDNA first-strand synthesis kit (Cat. No. KR211) according to a standard protocol (Tiangen Biotech Co., Ltd., Beijing, China). RT–qPCR was performed via a miRcute plus miRNA qPCR kit (SYBR Green) (Tiangen Biotech Co., Ltd., Beijing, China) on an ABI 7500 Real-Time PCR System (Applied Biosystems, Foster City, USA). The relative levels of individual miRNAs were calculated after normalization to the levels of miR16 and let7a in the corresponding sample. The specific primers used for RT–qPCR were synthesized by Sangon Biotech (Shanghai) Co., Ltd., and are listed in Supplementary Table S3.

### Multivariate analysis to select miRNA markers

The AdaBoost model and support vector machine (SVM) model were utilized to complementarily evaluate the ability of miRNAs to discriminate between the HCC and non-HCC groups. Based on the AdaBoost algorithm with the R package “adabag”, the importance of miRNA *i* was defined as its weighted frequency presented in the model containing 100 weighted trees (cross-validation = 10). We randomly sampled 75% of the original samples 30 times to obtain the miRNAs’ average importance scores. Moreover, based on the linear SVM model, the importance of gene *i* could be measured as

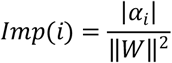

in which *W*= {*α*_*i*_}is the coefficient vector perpendicular to the support vectors. Then, the miRNAs’ average importance scores were derived by randomly sampling 75% of the original samples 500 times.

With the two definitions of importance above, the miRNAs with the minimum importance score were deleted sequentially from the DEM combination. The FDR, defined as the ratio of the prediction error rate of the original data to that of the label permutated data, was used to evaluate the miRNA combinations’ predictive effect and determine the optimal miRNA panel.

### Prediction modelling

The Z scores of the original data were randomly divided into a training set and a testing set at a ratio of 7:3. We compared the performances and stabilities of nine common models in the sklearn package by bootstrap sampling the training set 100 times. Then, the predictive capabilities of two models with relatively high area under the receiver operating characteristic curve (ROC) curve (AUC) values and low standard deviations were evaluated on the testing set. Furthermore, the signature panel was validated in a subsequent study.

SVC models were constructed with the combination of alpha-fetoprotein (AFP) and miRNA signature panels and with either AFP or miRNA signature panels. To compare their performance, we fit the mean of the ROC curves and calculated the 95% confidence interval using the “roc-utils” package, with 100 bootstrap samplings.

### Statistical analysis

The present study used GraphPad Prism 8.0 (GraphPad Software, Inc., San Diego, USA) for performing statistical calculations and data plotting. The differences between two independent samples in Fig. 2 and Fig. 3 were evaluated using two-tailed Student’ s t tests. Since the expression of miRNA was not normally distributed according to the Shapiro–Wilk test, the DEMs were evaluated using the Mann–Whitney test, the results of which are shown in Fig. 4, Fig. 5, and Fig. 6. All the tests were two-tailed unless otherwise indicated. We considered a P value ≤ 0.05 to be statistically significant. Significance values were set as follows: ns (not significant), P > 0.05; *, P < 0.05; **, P < 0.01; and ***, P < 0.001.

## Results

### Fucosylated EV enrichment based on the GlyExo-Capture method

To capture fucosylated EVs, a GlyExo-Capture method was developed by immobilizing LCA on hydroxyl macromolecular magnetic complexes. The fucosylated EVs of a maximum of 96 samples were isolated simultaneously by an EV automatic isolation and purification system (Hotgen Biotech Co., Ltd., Beijing, China). The reagents were prepackaged in a 96-well plate, and the whole isolation process took approximately 11 min. The workflow describing the GlyExo-Capture method is summarized in Fig. 1, and detailed information is listed in Supplementary Table S4.

**Figure 1.**
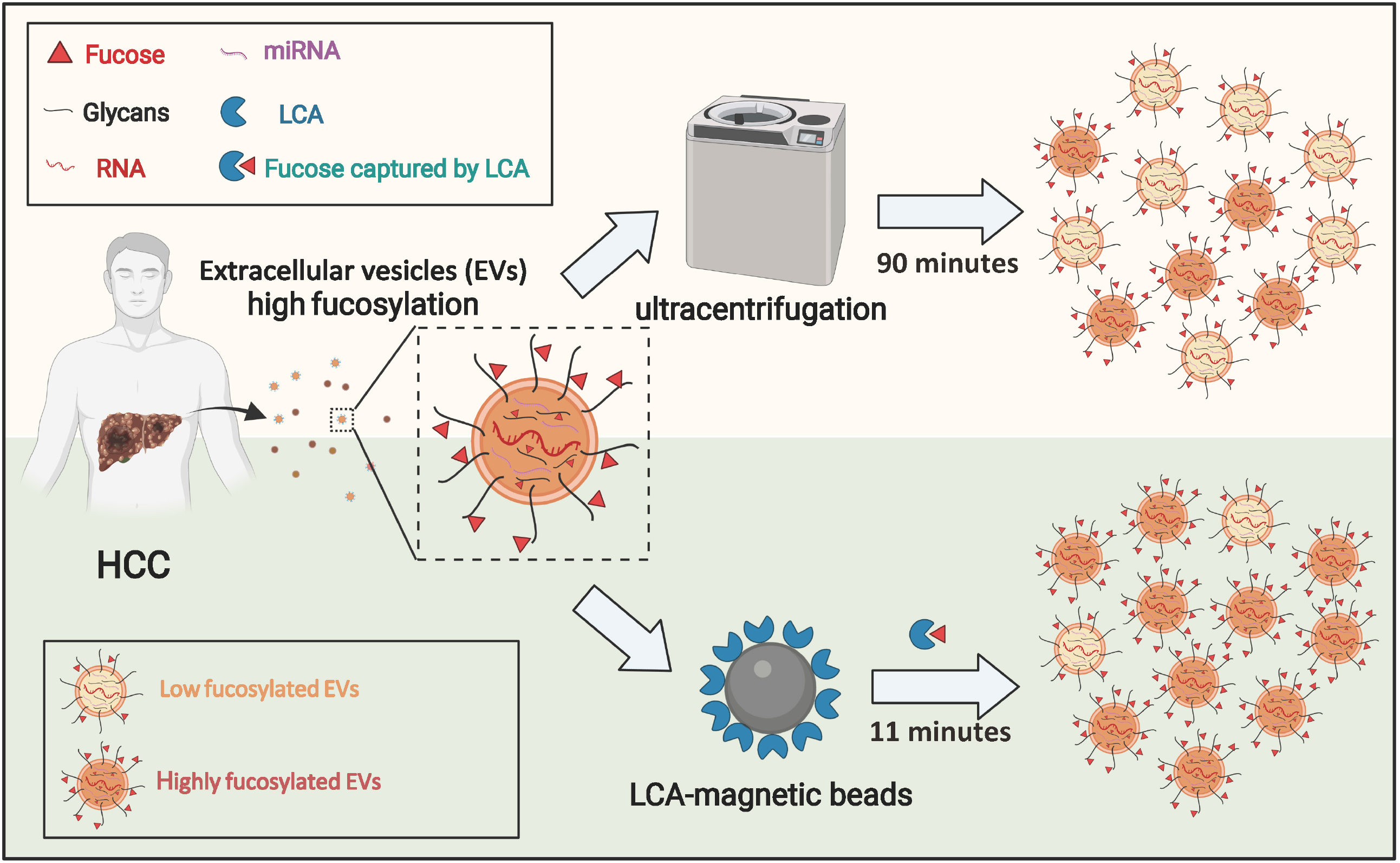
Workflow of the GlyExo-Capture technique. LCA magnetic beads can specifically capture fucosylated EVs from serum samples and cell supernatants. The whole process of preparing 96 samples takes only 11 min. LCA: Lens culinaris lectin, HCC: hepatocellular carcinoma.

To compare the morphology of EVs isolated by the different methods, the EVs from cell supernatant were characterized. The EVs isolated by the differential ultracentrifugation method (UC-EVs) were used as controls. TEM analysis showed that the fucosylated EVs and UC-EVs were oval or bowl shaped (Fig. 2a and b)). The average size of the fucosylated EVs (106.5±6.7 nm) was smaller than the average size of the UC-EVs (137.5±0.4 nm), but the fucosylated EVs had a broader size distribution (Fig. 2a and b). Immunoblot and ExoView analysis showed the expression of tetraspanins, i.e., TSG101, and ALIX (Fig. 2c and d), indicating the successful isolation of fucosylated EVs from the serum samples. Notably, increased staining intensity of tetraspanins was observed in the fucosylated EVs compared to the UC-EVs when the same doses of EVs were used, demonstrating that higher EV purification was achieved when this GlyExo-Capture method was used (Supplementary Fig. S1).

**Figure 2.**
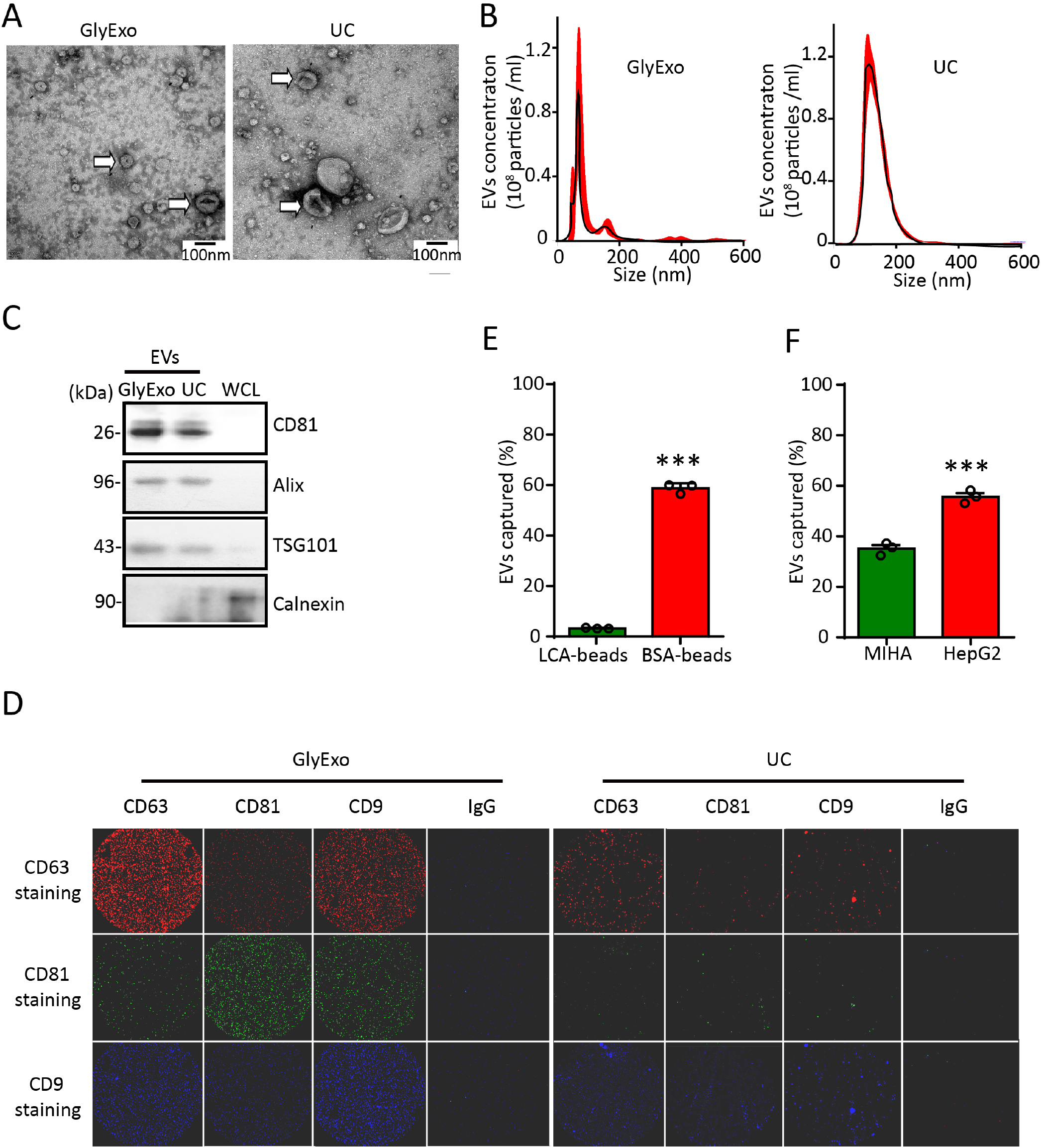
Characterization of fucosylated EVs isolated by the GlyExo-Capture technique. (a, b) EVs isolated by the GlyExo-Capture technique (GlyExo) and differential ultracentrifugation (UC) were analysed via TEM (a) or NTA (b). The representative EVs are highlighted by arrows in (a). Scale bars, 100 nm. (c) GlyExo-EVs, UC-EVs, and whole-cell lysates (WCLs) were analysed by western blotting using the indicated antibodies. (d) GlyExo-EVs and UC-EVs were analysed by ExoView using the indicated antibodies. (e) The UC-EVs of HepG2 cells were incubated with LCA-immobilized beads or BSA-immobilized beads, and the bound fractions were subjected to quantitative analysis. The doses of EVs were quantified using a Qubit Protein Assay and a Qubit 3.0 fluorometer. The amount of EVs captured was calculated as the bound fraction/UC-EVs loaded. (f) The UC-EVs of HepG2 or MIHA cells were incubated together with LCA-immobilized beads, and the bound fraction was subjected to quantitative analysis. Cell-based studies were performed independently at least three times, each yielding comparable results. The differences were evaluated using two-tailed Student’ s t tests. *, P < 0.05; **, P < 0.01; ***, P < 0.001.

To estimate the binding efficiency of LCA-immobilized beads, the UC-EVs were incubated with LCA-immobilized beads or BSA-immobilized beads, and the unbound fractions were subjected to quantitative analysis. The analysis showed that incubation of the UC-EVs with LCA-immobilized beads but not with BSA-immobilized beads resulted in a significant decrease (by 40%) in UC-EV particle numbers in the unbound fraction, suggesting that approximately 60% of HepG2-derived EVs were captured by LCA-immobilized beads (Fig. 2e). Specifically, the proportion of fucosylated EVs secreted by MIHA cells, which constitute a line of nontumorigenic hepatocyte cells, was lower than 40% (Fig. 2f). Therefore, the GlyExo-Capture technique is a fast and reliable method to isolate fucosylated EVs from cell supernatants and human sera in a high-throughput manner.

### Cellular uptake of fucosylated EVs

To characterize fucosylated EVs, the uptake patterns of the fucosylated EVs by their parent cells were first evaluated. To this end, the EVs collected by the GlyExo-Capture beads were identified as enriched or HF-EVs, and the EVs in the unbound supernatant subjected to differential ultracentrifugation were identified as LF-EVs. These EVs were labelled with the fluorescent dye DiD, and the uptake behaviours were analysed by flow cytometry. The results revealed that the HF-EVs exhibited time- and dose-dependent uptake (Fig. 3a and b). Notably, the HF-EVs exhibited significantly higher uptake efficiency than did the LF-EVs (Fig. 3c and d). Notably, the enzymatic removal of N-glycans through PNGase F reduced the enhanced uptake efficiency of HF-EVs to levels comparable to those of LF-EVs (Fig. 3c and d). Thus, the glycosylated form of EVs may be responsible for their adherence and internalization.

**Figure 3.**
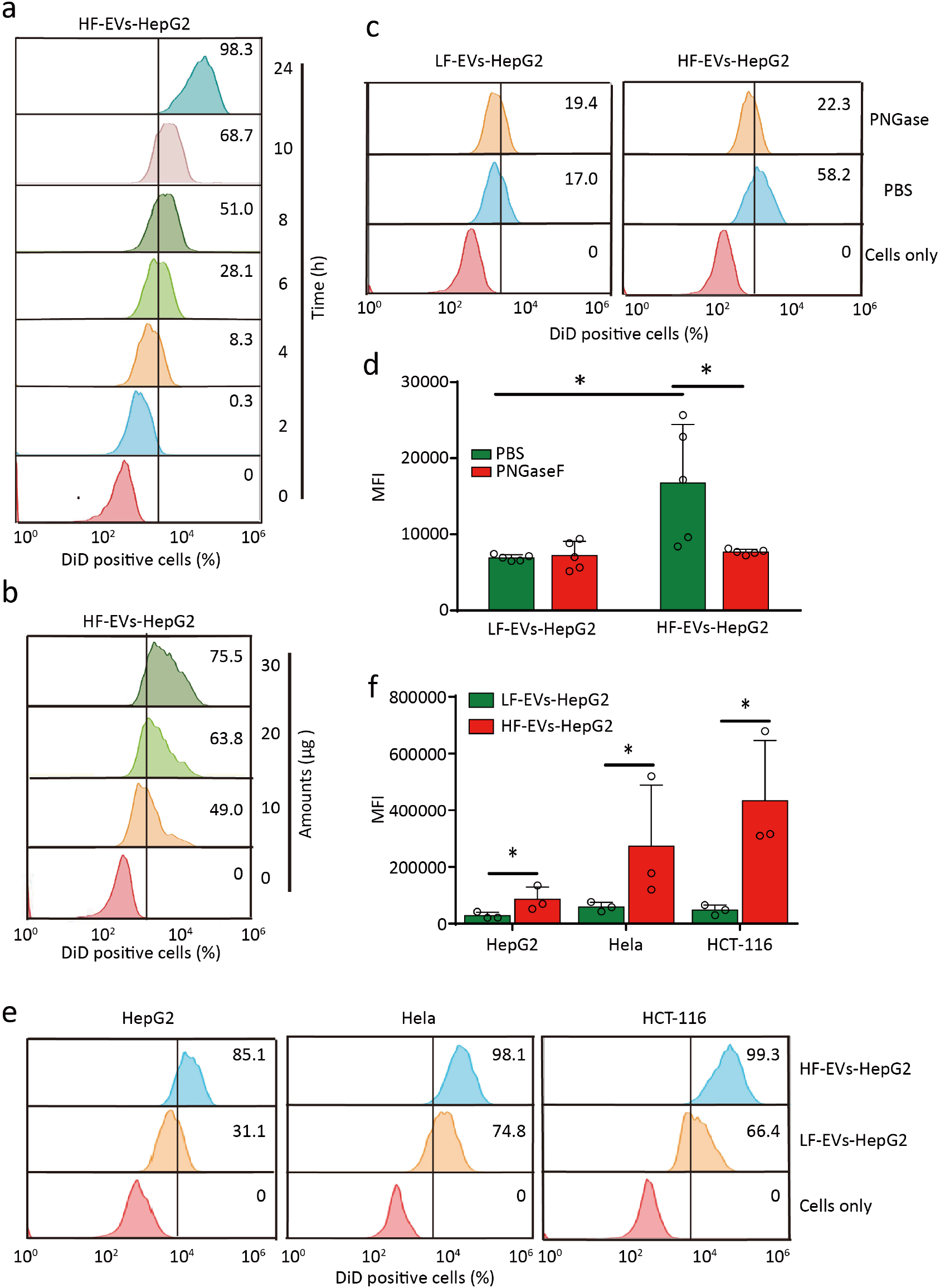
Cellular uptake of fucosylated EVs. (a, b) Flow cytometry analysis of DiD-labelled HF-EV uptake by parental HepG2 cells. Afterwards, 20 μg of DiD-labelled HF-EVs were added to HepG2 cells for the indicated time (a), or different amounts of DiD-labelled HF-EVs were added to HepG2 cells for 4 hours (b). (c, d) Flow cytometry analysis of DiD-labelled HF-EV or LF-EV uptake by parental HepG2 cells in the presence or absence of PNGase F. The intracellular fluorescence intensity from DiD-labelled EVs was quantitated as the percentage of positive cells (c) and the mean fluorescence intensity (MFI; d). (e, f) Flow cytometry analysis of DiD-labelled HF-EV or LF-EV uptake by HepG2, HCT-116, and HeLa cells. The intracellular fluorescence intensity from DiD-labelled EVs was quantitated as the percentage of positive cells (e) and the MFI (f). The cell-based studies were performed independently at least three times with comparable results. The differences were evaluated using two-tailed Student’ s t tests. *, P < 0.05; **, P < 0.01; ***, P < 0.001.

After demonstrating the improved uptake of HF-EVs by their parent cells, we further evaluated whether this phenomenon was cell specific. To this end, LF-EVs and HF-EVs labelled with DiD released from HepG2 cells were exposed to various cells. Consistent with the above results, HepG2 HF-EVs were also quickly loaded by HepG2 cells (Fig. 3e and f). However, HCT-116 and HeLa cells accumulated significantly less HF-EVs-HepG2 than LF-EVs-HepG2, which indicates a specific targeting behaviour of fucosylated EVs (Fig. 3e and f).

### miRNA profiles of fucosylated EVs in HCCs

As glycans are highly expressed on cancer EVs, the finding of enriched fucosylated EV miRNA cargos may hold tremendous potential to identify more specific and sensitive cancer EV biomarkers. To this end, a cohort of 61 patients with HCC, 24 patients with hepatitis and 30 healthy subjects were recruited. NGS was used to determine the global profile of the fucosylated EV miRNA cargos. For each sample, at least 15 M raw reads were generated, and a total of 1938 known miRNAs were identified. Patients with hepatitis and healthy subjects were grouped together as non-HCC controls unless otherwise stated. According to the results of the DEM (FC>2, *p*<0.05) analysis, 52 miRNAs were upregulated and 23 downregulated in the HCC group compared with the non-HCC group (Supplementary Fig. S2; Supplementary Table S5). LDA based on an expression matrix of 75 DEMs showed that three distinct clusters generally occurred: an HCC group, a hepatitis group, and a healthy group (Fig. 4a). Compared with DEMs in The Cancer Genome Atlas (TCGA) dataset, 19 DEMs of fucosylated EVs showed the same expression profiles (Fig. 4b). Notably, the targeted mRNAs were mainly enriched in pathways associated with cancer, endocytosis, Ras signalling, Rap1 signalling, and proteoglycans in cancer (Supplementary Fig. S3).

**Figure 4.**
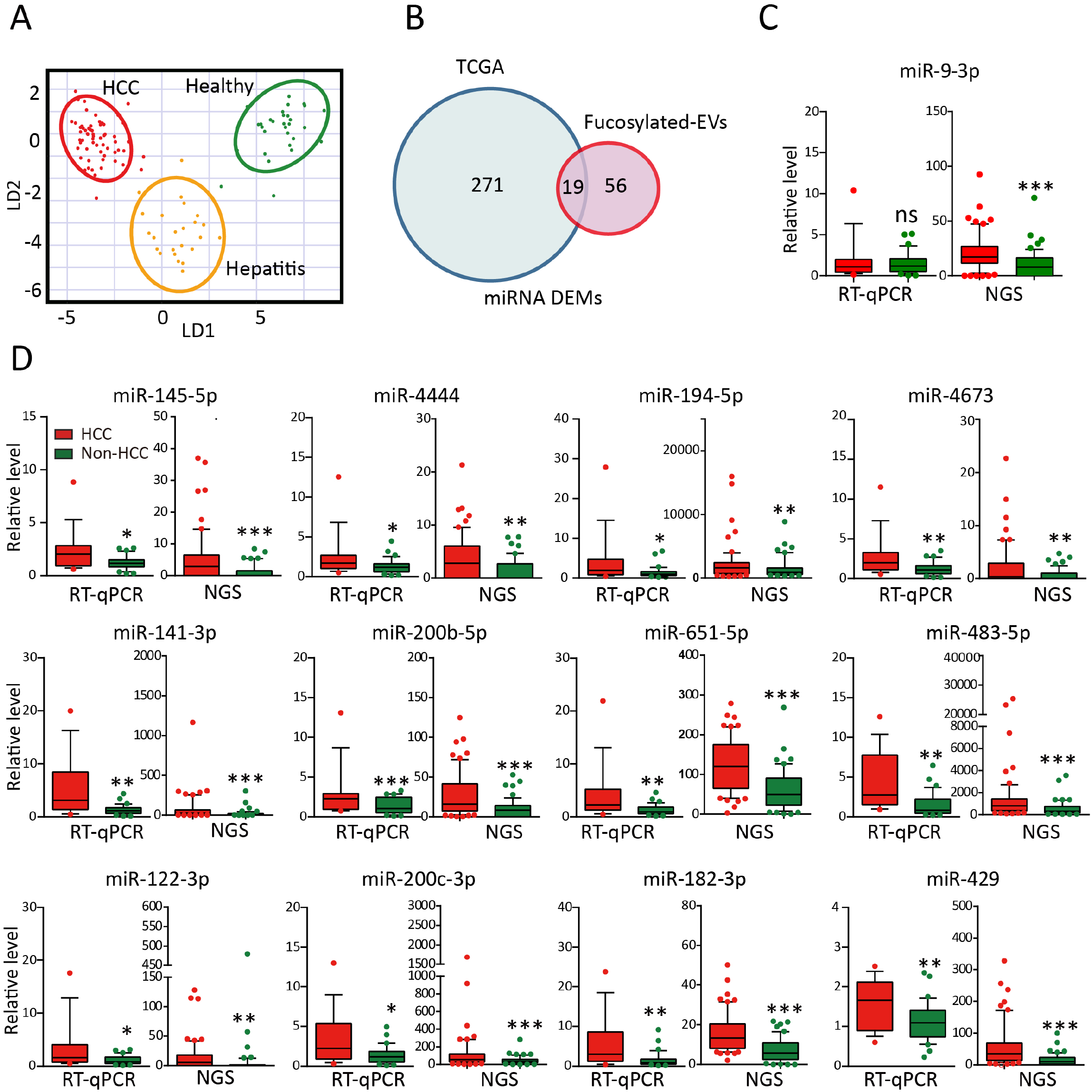
Verification of fucosylated EV miRNA expression in HCCs. (a) LDA based on an expression matrix of the 75 DEMs identified by NGS. The red dots represent patients with HCC, the yellow dots represent patients with hepatitis, and the green dots represent healthy controls. (b) Venn diagram showing the DEMs shared between the TCGA dataset (blue circles) and the fucosylated EV miRNA dataset (red circles). (c, d) Expression of 13 randomly selected DEMs according to RT–qPCR or NGS. The expression of the miRNA levels was normalized to those in the non-HCC group (RT–qPCR). The expression pattern of miR-9-3p is shown in (c). The other 12 miRNAs are shown in (d). The differences were evaluated using the Mann–Whitney test. *, P < 0.05; **, P < 0.01; ***, P < 0.001.

To verify the NGS data, another cohort of 48 participants (16 patients with HCC, 16 patients with hepatitis, and 16 healthy controls) was recruited, and 13 randomly selected DEMs were confirmed by RT–qPCR analysis. Except for 1 DEM, 12 out of 13 DEMs exhibited expression patterns similar to those detected on the basis of the NGS data (Fig. 4c and d).

### Identification of HCC-specific miRNA signatures in fucosylated EVs

Using multivariate analysis, we selected a panel of 6 miRNAs from the 75 DEMs based on the NGS results (Fig. 5a). The RT–qPCR results showed that miR-215-5p and miR-432-5p expression was either downregulated or unchanged in the HCC group (Fig. 5b), whereas miR-483-5p, miR-182-3p, miR-651-5p, and miR-429 were all significantly upregulated in the HCC group (Fig. 5c). The expression trend of these 4 miRNAs was consistent between the RT–qPCR and NGS data (Fig. 5c). Notably, these 4 miRNAs were weakly correlated with each other, which implied their high complementary feature in diagnostic use (Fig. 5d). Therefore, the 4-miRNA panel from fucosylated EVs was chosen as a potential signature biomarker for further validation.

**Figure 5.**
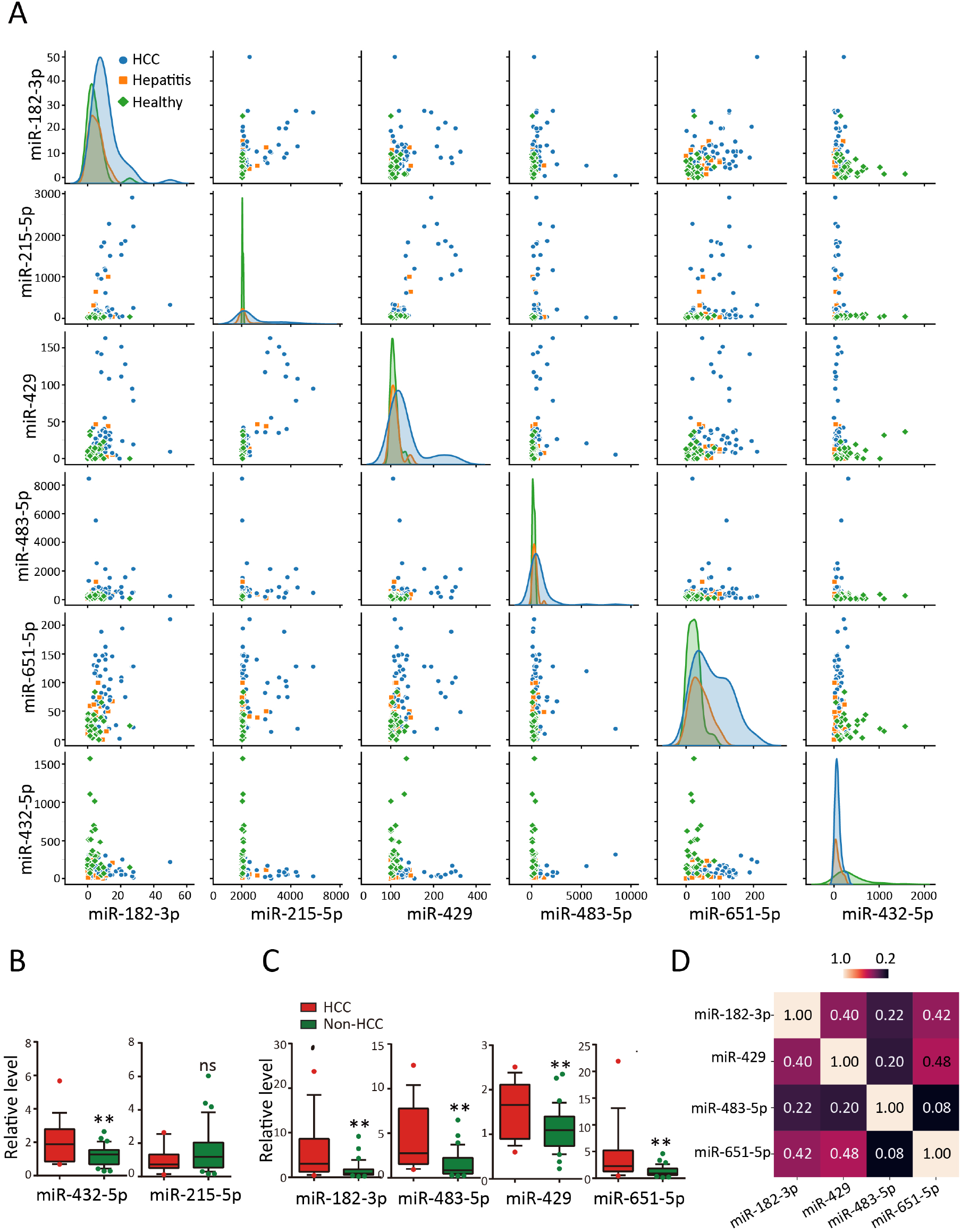
Identification of HCC-specific miRNA signatures in fucosylated EVs. (a) Paired scatterplot matrix of 6 miRNA markers selected on the basis of the results of a multivariate analysis of the original samples. (b, c) Expression of miR-432-5p and miR-215-5p (b) and miR-183-3p, miR-483-5p, miR-429, and miR-651-5p (c) according to RT–qPCR of the HCC and non-HCC control groups. (d) Spearman correlation heatmap assay of the 4 miRNAs (miR-183-3p, miR-483-5p, miR-429, and miR-651-5p). The differences were evaluated using the Mann–Whitney test. *, P < 0.05; **, P < 0.01; ***, P < 0.001.

### Diagnostic evaluation of the 4-miRNA signature in fucosylated EVs

Based on the 4-miRNA signature in fucosylated EVs, different prediction models, including gradient boosting, AdaBoost, random forest, bagging, decision tree, Gaussian NB, K-neighbours, SVC, and logistic regression, were employed. The cohort comprising 61 patients with HCC, 24 patients with hepatitis, and 30 healthy controls was randomly divided into a training sample set and a testing sample set at a ratio of approximately 7:3. Compared with the other models, the AdaBoost model and random forest model achieved higher AUC values with lower standard deviations (Fig. 6(a)). Therefore, these two models were generated to detect HCC samples. As shown, the ROC analysis of the AdaBoost model showed that the 4-miRNA-based screening signature yielded an AUC of 0.86 for the detection of HCC from non-HCC controls in the testing sample (Figure 6(b)).

**Figure 6.**
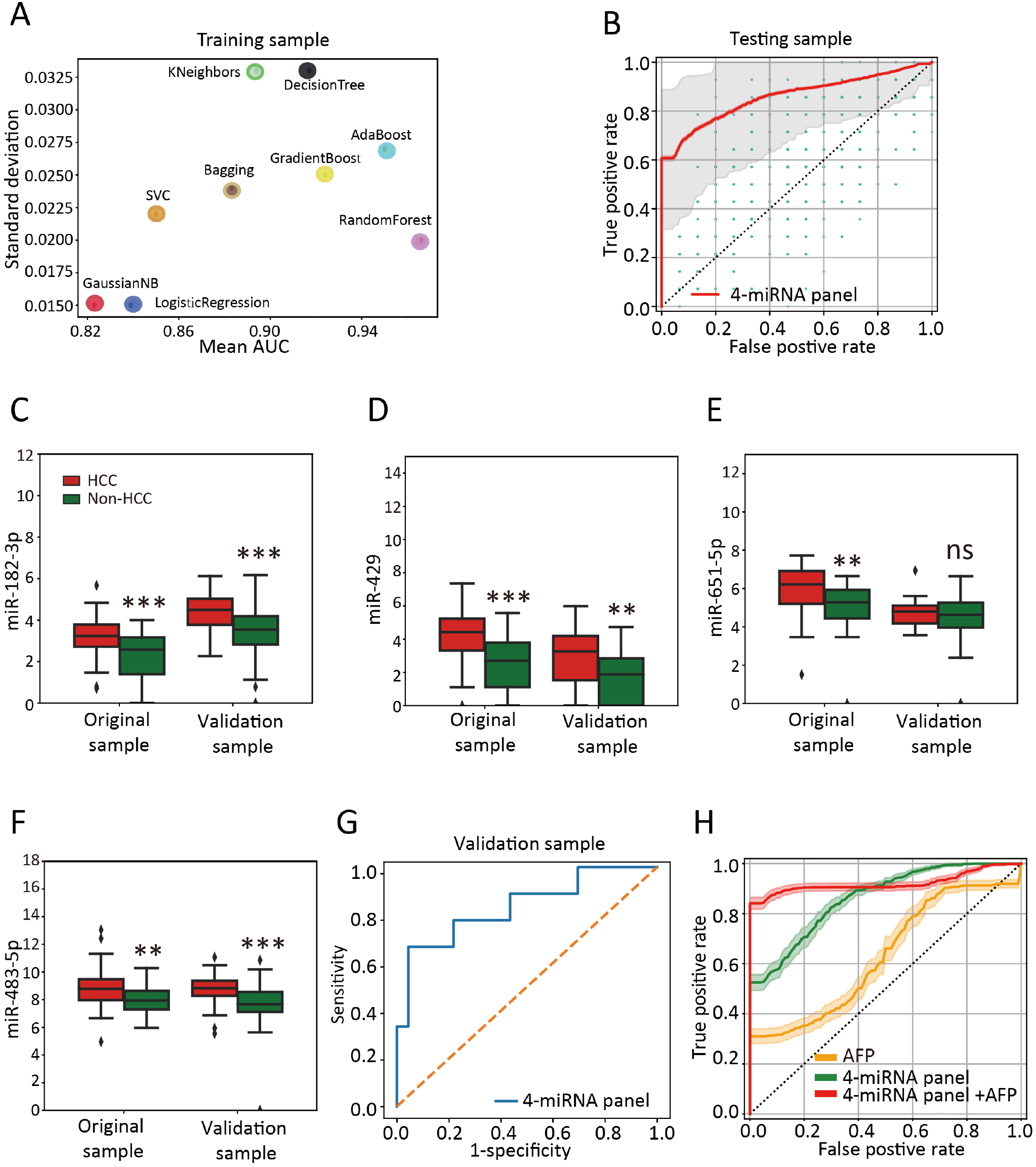
Diagnostic evaluation of the 4-miRNA panel in fucosylated EVs. (a) Prediction efficacies of widely used models, including gradient boosting, AdaBoost, random forest, bagging, decision tree, Gaussian NB, K-neighbours, SVC, and logistic regression, based on the 4-miRNA panel in the training sample. (b) Prediction capabilities according to the AdaBoost model based on the 4-miRNA panel (red line, AUC=0.86) or 4 randomly selected miRNAs in the testing sample (blue dots). (c-f) Expression of miR-182-3p (c), miR-429 (d), miR-651-5p (e), and miR-483-5p (f) according to NGS data from the original sample and the validation sample. (g) Verification of the 4-miRNA panel as a biomarker of HCC in the validation sample (AUC=0.84). (h) The 4-miRNA panel with AFP improved the HCC diagnostic accuracy (AUC=0.92). The differences were evaluated using the Mann–Whitney test. *, P < 0.05; **, P < 0.01; ***, P < 0.001.

We next monitored whether the 4-miRNA-based screening signature can discriminate HCC from non-HCC controls in another validation sample set composed of 30 patients with HCC, 52 patients with hepatitis, and 23 healthy controls. Specifically, using RT–qPCR, we first evaluated the expression profiles of the 4-miRNA signature in the original samples and the validation samples. As expected, similar expression trends were observed between the two sample sets (Figure 6(c-f)). Notably, the 4-miRNA-based diagnostic signature also performed well at predicting HCC (AUC: 0.84; Figure 6(g)). The 4-miRNA signature was independent of AFP, and a combined model with AFP yielded an increased AUC – 0.92 (Figure 6(h)).

## Discussion

In clinical practice, many glycoproteins or glycoconjugates with altered glycosylation have been used as serological markers to diagnose cancers (17,18). Cancer EVs are rich in many aberrant glyco-signatures, such as mannose, complex type N-glycans, polylactosamine, and sialylated glycans (11). To date, several strategies have been developed to characterize the glycan profiles of EVs (7). However, efficient and high-throughput isolation methods are still technically challenging. In this study, we developed a novel GlyExo-Capture technique to isolate EVs from serum samples and a panel of cancer cells using the fucose-specific LCA. Compared with EVs isolated by ultracentrifugation, EVs captured by this GlyExo-Capture technique were morphologically intact with a broader size distribution pattern and increased purity, supporting the advantage of the glyco-lectin affinity method over ultracentrifugation. The GlyExo-Capture technique is aimed toward glycan specificity rather than size specificity. Therefore, this technique could capture not only large vesicles but also smaller vesicles. Moreover, the whole process of preparing 96 samples took only 11 min.

Reports have revealed the role and impact of glycosylation in EV biodistribution and uptake (19). Cleavage of terminal sialic acid residues occurs more readily in EVs in the lungs and axillary lymph nodes than in nontreated EVs (20). For most of the cell lines tested, desialylated EVs presented increased uptake efficiency (21). In our study, we found that the fucosylation of EVs strongly increased their uptake efficiency in parent cells but not in the other cancer cell lines tested. Specifically, cleavage of EV N-glycans by PNGase F digestion abolished the increase in the uptake of fucosylated EVs. The different roles of sialic acid and fucose in the interaction between EVs and recipient cells need further investigation. Nevertheless, interestingly, our results suggest that EVs and cell fucosylation can impact EV uptake and, therefore, may drive specific EV organ biodistribution.

One of the most critical aspects of research concerning glycosylated EVs involves analysing their nucleic acid cargo, particularly small RNAs. Circulating miRNA biomarkers in the blood have been widely applied to disease diagnosis. Previous studies have indicated that miRNA biomarkers can be enriched in EVs (22,23). The establishment of the GlyExo-Capture method makes the detection of serum fucosylated EV-derived miRNA biomarkers a reality. We isolated fucosylated EVs from serum samples of patients with HCC and subjects without HCC by the GlyExo-Capture technique. Analysis of serum fucosylated EV-derived miRNAs between the HCC and non-HCC groups revealed a unique miRNA expression profile in the HCC group. Among the 75 DEMs from fucosylated EVs, 19 were also identified in the HCC dataset from TCGA, indicating that the miRNAs in the enriched fucosylated EV fractions could reflect the miRNA profile in the primary tumour site and may be involved in critical miRNA functions in cancer progression. In particular, a panel of 4 miRNAs was chosen as a signature for HCC detection. Based on the samples in this study, the 4-miRNA panel shows great performance and, in combination with AFP, significantly increases HCC detection efficiency. The potential application of this GlyExo-Capture technique for pancancer detection warrants further investigation.

In summary, the GlyExo-Capture technique, which takes advantage of the ability of lectins to specifically select various types of glycans, can effectively capture and enrich sugar-coated EVs. This GlyExo-Capture technique is user friendly, fully automatic and robust. The high-throughput feature supports the usage of this technique in tumour liquid biopsy and may open new avenues to determine the roles of EV glycosylation in cancer biology and therapy.

## Acknowledgments

We appreciate the support and participation of the physicians and patients in this study. We thank Prof. Zhong for editing the manuscript. We also thank Yufei Wang and Wade Wang (Quantum Design China & NanoView Biosciences) for supporting the characterization of extracellular vesicles. A graphical abstract was created and exported with BioRender.com under a paid subscription.

## Figure legends

**Supplementary Figure S1.**
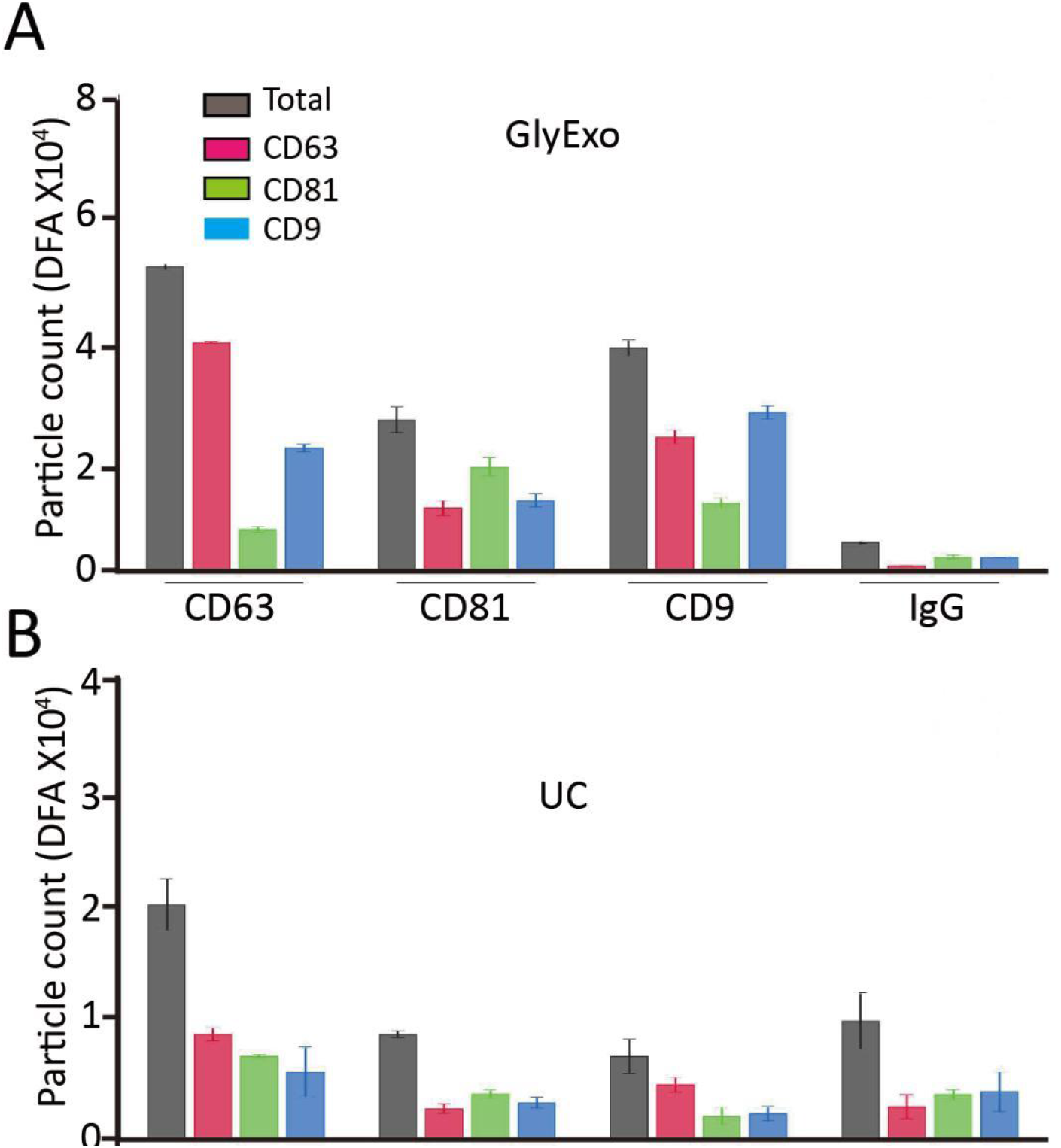
Characterization of fucosylated EVs, in relation to Figure 2. A-B. Histogram of the results of an ExoView analysis of GlyExo-EVs (A) and UC-EVs (B) using the indicated antibodies. IgG was used as a negative control. DFA: Dilution factor adjusted.

**Supplementary Figure S2.**
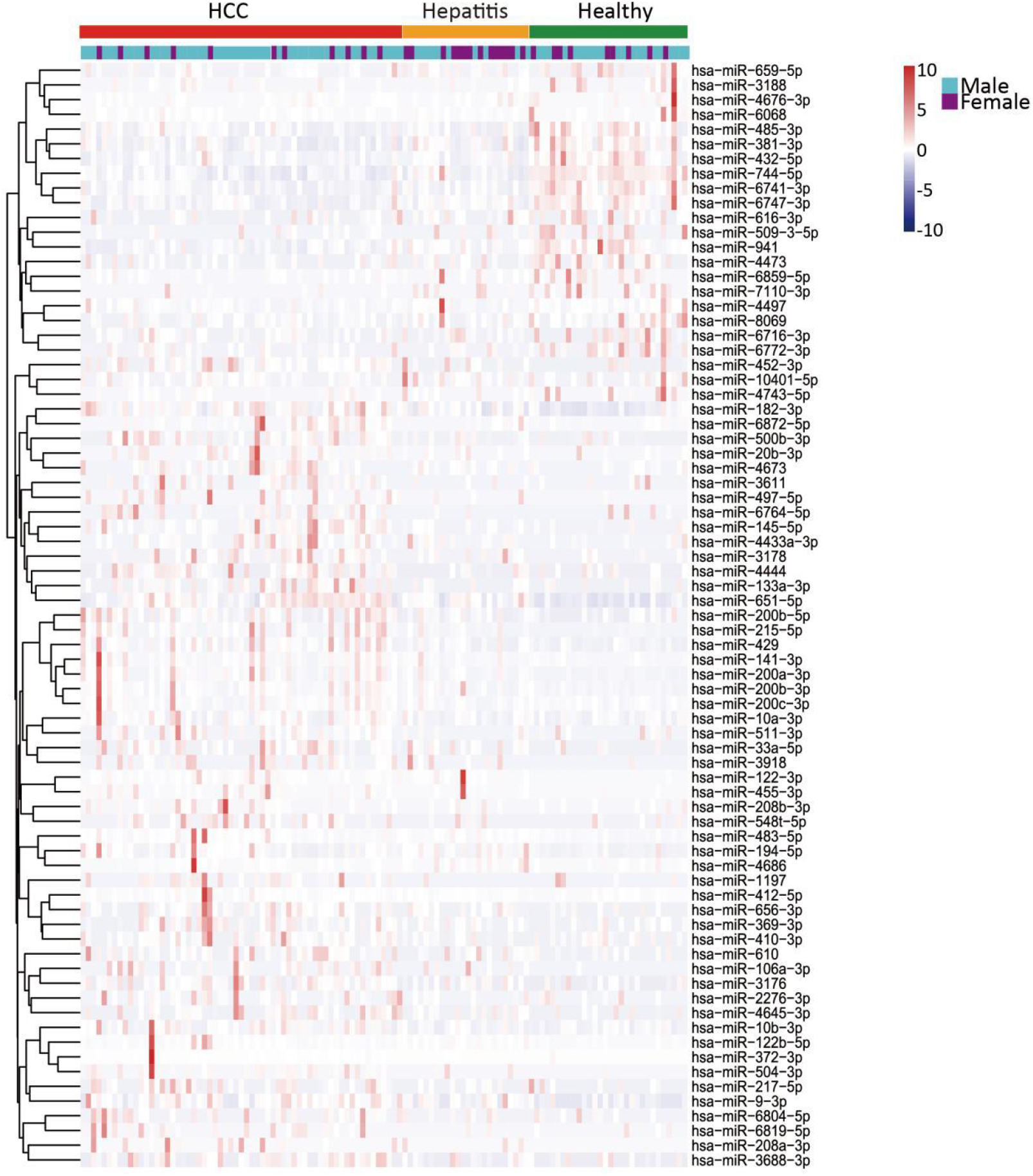
miRNA profiles of fucosylated EVs in HCC, in relation to Figure 4. Heatmap of the expression levels of 75 DEMs across all 115 samples in the fucosylated EV-derived miRNA dataset.

**Supplementary Figure S3.**
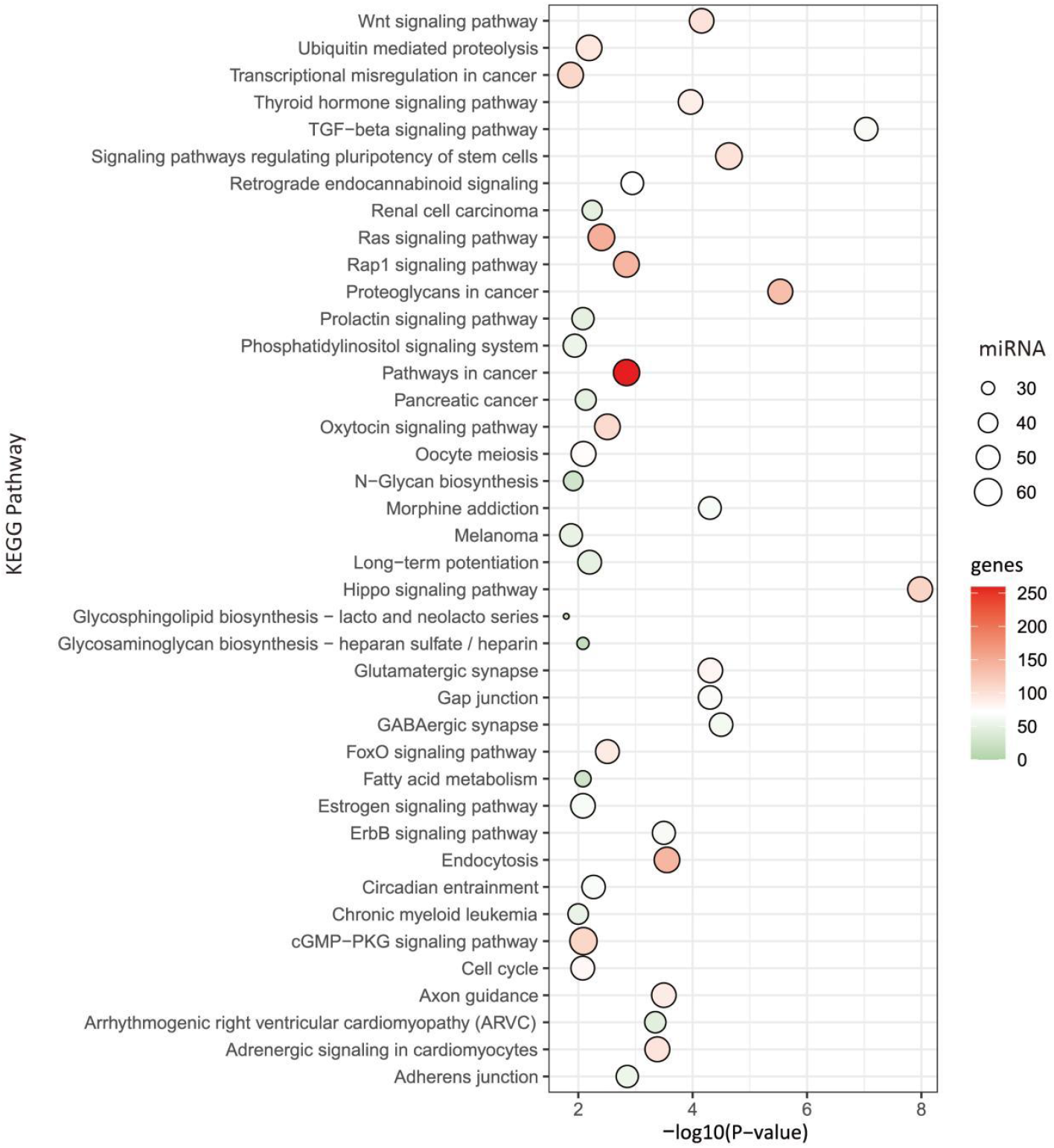
miRNA pathway analysis of fucosylated EVs in HCC, in relation to Figure 4. Kyoto Encyclopedia of Genes and Genomes (KEGG) enrichment analysis (top 40 terms ranked by P value) of mRNAs targeted by the 75 DEMs identified in the fucosylated EV-derived miRNA dataset.

**Supplementary Table S1.**
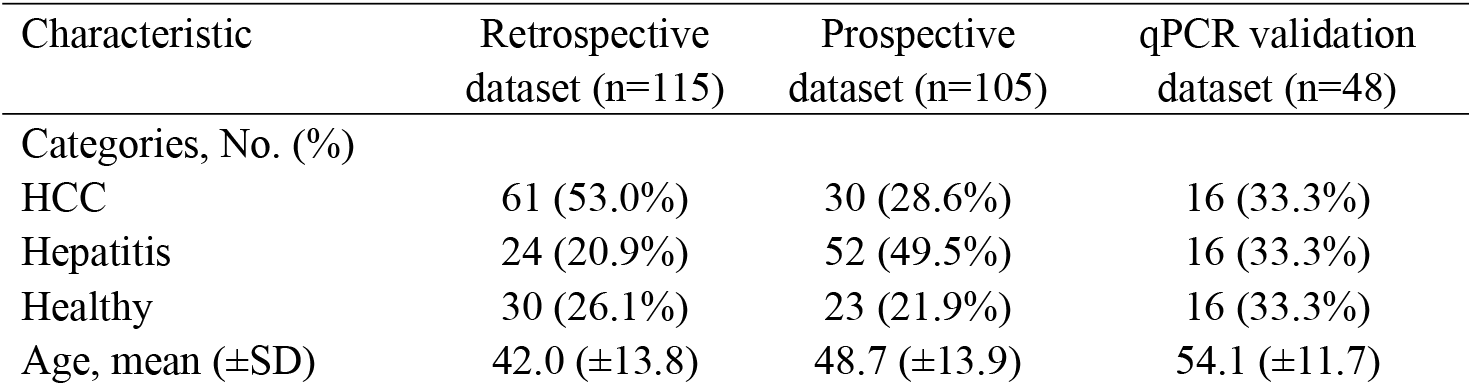
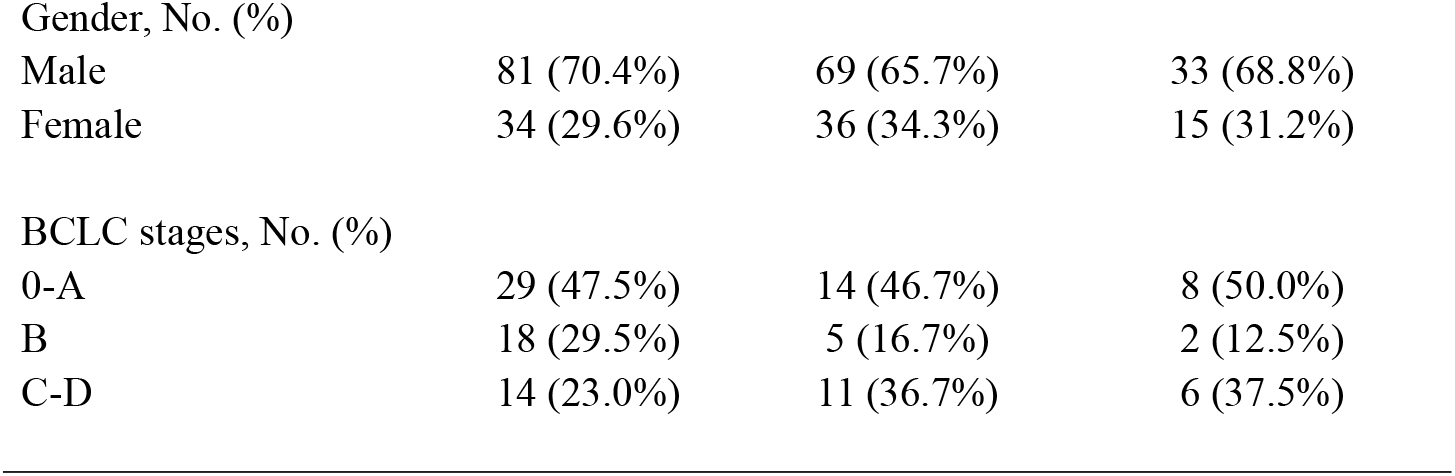
The demographic and clinicopathologic characteristics of the patients in three datasets

| Characteristic | Retrospective dataset (n=115) | Prospective dataset (n=105) | qPCR validation dataset (n=48) |
| --- | --- | --- | --- |
| Categories, No. (%) |  |  |  |
| HCC | 61 (53.0%) | 30 (28.6%) | 16 (33.3%) |
| Hepatitis | 24 (20.9%) | 52 (49.5%) | 16 (33.3%) |
| Healthy | 30 (26.1%) | 23 (21.9%) | 16 (33.3%) |
| Age, mean (±SD) | 42.0 (±13.8) | 48.7 (±13.9) | 54.1 (±11.7) |
| Gender, No. (%) |  |  |  |
| Male | 81 (70.4%) | 69 (65.7%) | 33 (68.8%) |
| Female | 34 (29.6%) | 36 (34.3%) | 15 (31.2%) |
| BCLC stages, No. (%) |  |  |  |
| 0-A | 29 (47.5%) | 14 (46.7%) | 8 (50.0%) |
| B | 18 (29.5%) | 5 (16.7%) | 2 (12.5%) |
| C-D | 14 (23.0%) | 11 (36.7%) | 6 (37.5%) |

**Supplementary Table S2.**
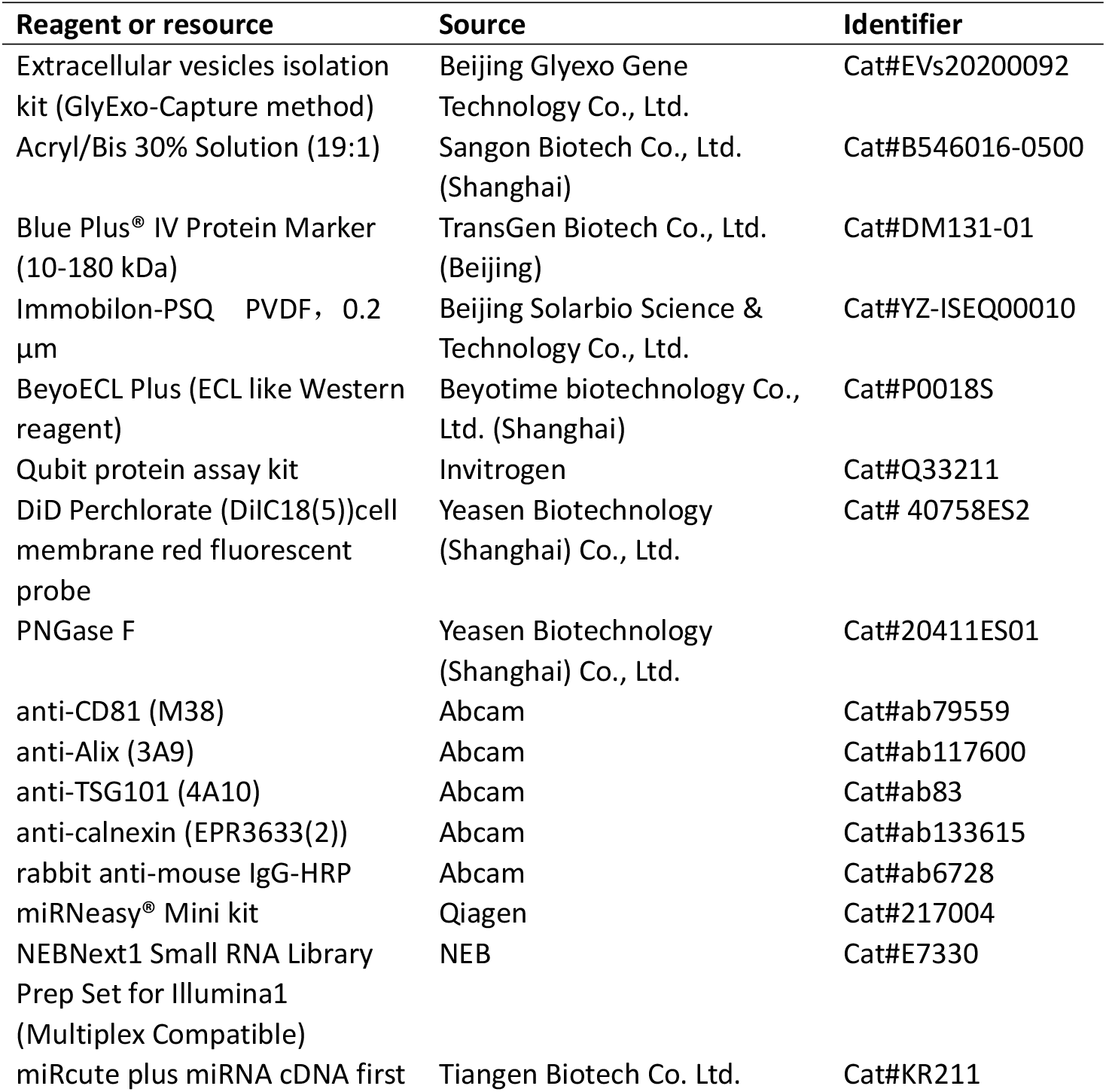
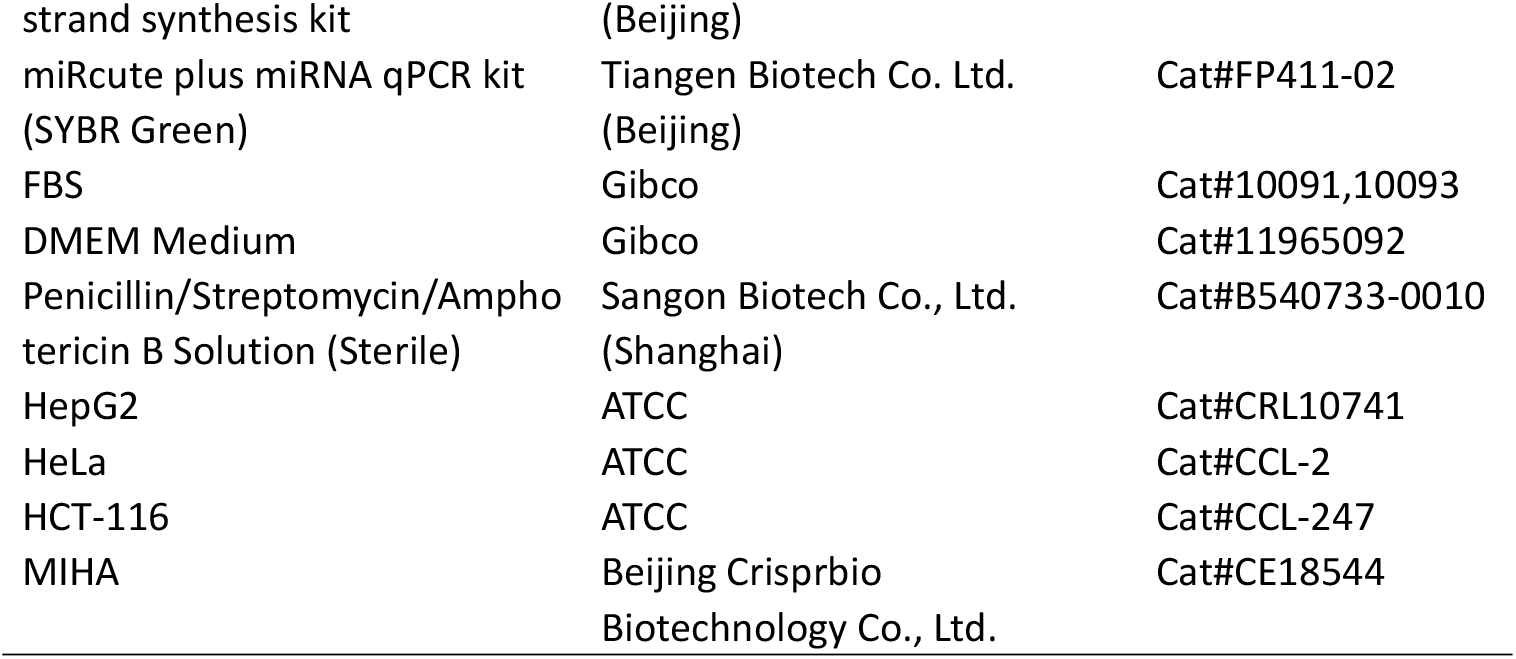
Key resources table

**Supplementary Table S3.**
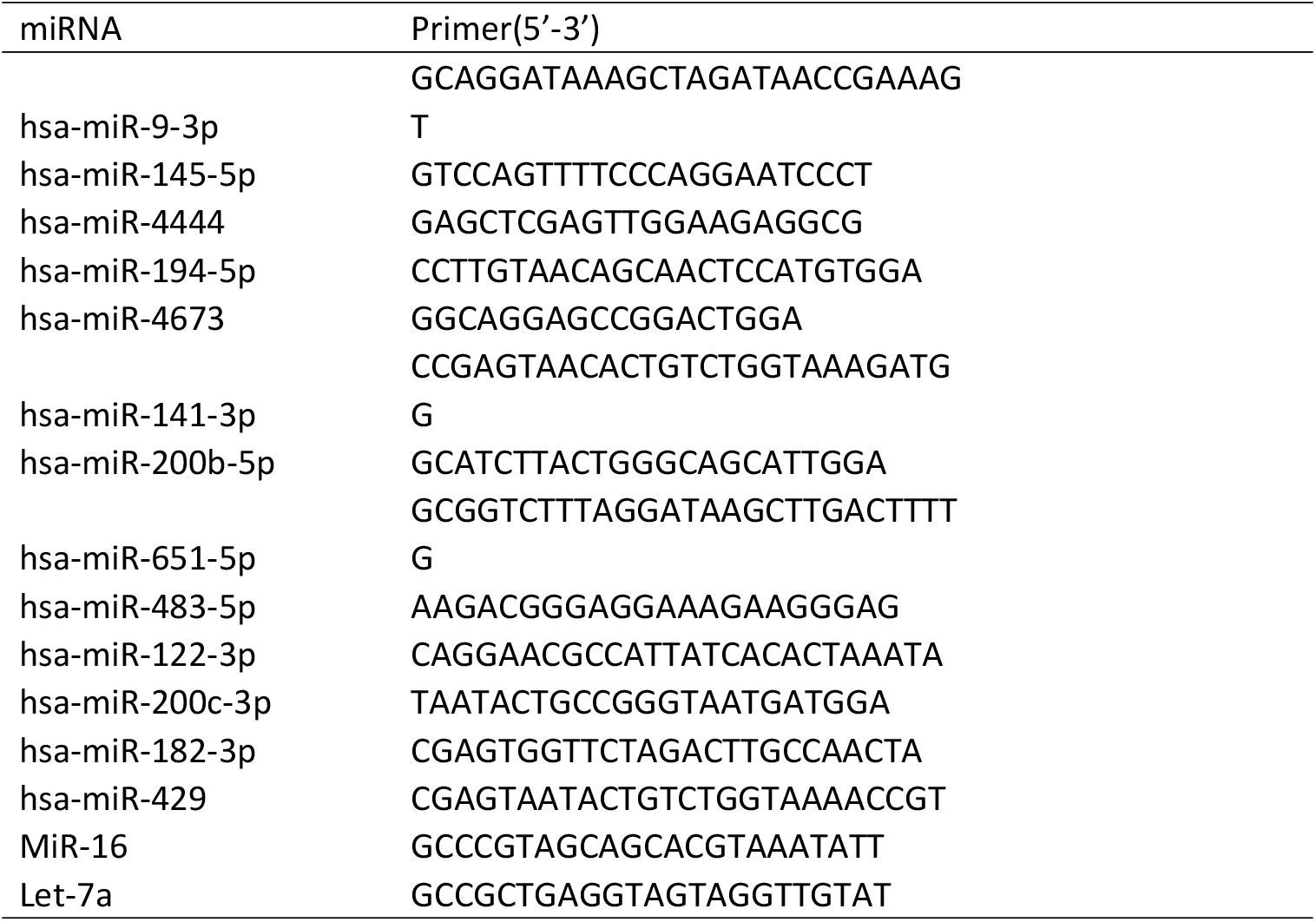
The primers used for qRT–PCR

**Supplementary Table S4.** The procedure for isolating EVs by GlyExo-Caputre technique

| Step | Well | Volume (μL) | Mixing speed | Mixing time (minutes) | Precipitation time (seconds) | Magnetic absorption times |
| --- | --- | --- | --- | --- | --- | --- |
| MIX | 1 | 750 | LOW | 1 | 240 | 1 |
| WASH | 2 | 600 | LOW | 1 | 1 | 1 |
| ELUTE | 3 | 250 | LOW | 1 | 240 | 1 |

**Supplementary Table S5.** The specificity and sensitivity of TOP 45 DEMs in identifying HCC patients in our serum fucosylated-EVs miRNA dataset ranked by p-adj value.

| miRNA name | log<sub>2</sub>FC | p-adj | sensitivity | specifity | Youden | AUC |
| --- | --- | --- | --- | --- | --- | --- |
| hsa-miR-744-5p | -1.09 | 1.45E-11 | 0.95 | 0.63 | 0.58 | 0.79 |
| hsa-miR-215-5p | 2.49 | 2.36E-11 | 0.56 | 0.93 | 0.48 | 0.78 |
| hsa-miR-200a-3p | 1.83 | 5.35E-11 | 0.74 | 0.72 | 0.46 | 0.77 |
| hsa-miR-429 | 1.90 | 1.31E-08 | 0.67 | 0.81 | 0.49 | 0.78 |
| hsa-miR-941 | -1.10 | 2.97E-08 | 0.77 | 0.70 | 0.47 | 0.78 |
| hsa-miR-483-5p | 1.68 | 3.90E-07 | 0.54 | 0.80 | 0.34 | 0.70 |
| hsa-miR-182-3p | 1.19 | 4.40E-07 | 0.85 | 0.61 | 0.46 | 0.80 |
| hsa-miR-6741-3p | -1.61 | 9.24E-07 | 0.89 | 0.59 | 0.48 | 0.74 |
| hsa-miR-10b-3p | 1.40 | 1.41E-06 | 0.51 | 0.89 | 0.40 | 0.73 |
| hsa-miR-432-5p | -1.25 | 1.85E-06 | 0.92 | 0.50 | 0.42 | 0.72 |
| hsa-miR-200c-3p | 1.44 | 9.48E-06 | 0.49 | 0.81 | 0.31 | 0.64 |
| hsa-miR-141-3p | 1.74 | 1.34E-05 | 0.51 | 0.81 | 0.32 | 0.65 |
| hsa-miR-651-5p | 1.03 | 1.48E-05 | 0.48 | 0.94 | 0.42 | 0.78 |
| hsa-miR-6747-3p | -1.22 | 5.19E-05 | 0.77 | 0.63 | 0.40 | 0.76 |
| hsa-miR-200b-3p | 1.31 | 5.52E-05 | 0.69 | 0.80 | 0.49 | 0.75 |
| hsa-miR-145-5p | 2.15 | 1.18E-04 | 0.67 | 0.76 | 0.43 | 0.73 |
| hsa-miR-200b-5p | 1.44 | 1.60E-04 | 0.52 | 0.81 | 0.34 | 0.72 |
| hsa-miR-381-3p | -1.02 | 1.61E-04 | 0.75 | 0.67 | 0.42 | 0.72 |
| hsa-miR-455-3p | 3.13 | 3.69E-04 | 0.34 | 0.96 | 0.31 | 0.65 |
| hsa-miR-10a-3p | 1.22 | 1.84E-03 | 0.61 | 0.70 | 0.31 | 0.68 |
| hsa-miR-8069 | -1.35 | 2.50E-03 | 0.75 | 0.52 | 0.27 | 0.63 |
| hsa-miR-122-3p | 2.21 | 4.36E-03 | 0.69 | 0.72 | 0.41 | 0.72 |
| hsa-miR-500b-3p | 1.50 | 4.85E-03 | 0.62 | 0.80 | 0.42 | 0.71 |
| hsa-miR-6859-5p | -1.99 | 5.49E-03 | 0.89 | 0.35 | 0.24 | 0.56 |
| hsa-miR-9-3p | 1.06 | 6.23E-03 | 0.82 | 0.63 | 0.45 | 0.73 |
| hsa-miR-4433a-3p | 1.78 | 8.27E-03 | 0.44 | 0.81 | 0.26 | 0.63 |
| hsa-miR-6872-5p | 1.81 | 8.64E-03 | 0.46 | 0.74 | 0.20 | 0.60 |
| hsa-miR-4444 | 1.40 | 9.79E-03 | 0.74 | 0.69 | 0.42 | 0.72 |
| hsa-miR-504-3p | 2.40 | 1.20E-02 | 0.44 | 0.93 | 0.37 | 0.68 |
| hsa-miR-4686 | -1.82 | 1.26E-02 | 0.90 | 0.30 | 0.20 | 0.56 |
| hsa-miR-3176 | 1.13 | 1.27E-02 | 0.74 | 0.63 | 0.37 | 0.70 |
| hsa-miR-485-3p | -1.08 | 1.29E-02 | 0.66 | 0.57 | 0.23 | 0.63 |
| hsa-miR-4497 | -1.08 | 1.46E-02 | 0.72 | 0.54 | 0.26 | 0.62 |
| hsa-miR-369-3p | 1.43 | 1.52E-02 | 0.36 | 0.91 | 0.27 | 0.63 |
| hsa-miR-412-5p | 1.26 | 1.52E-02 | 0.70 | 0.54 | 0.24 | 0.60 |
| hsa-miR-4673 | 1.55 | 1.74E-02 | 0.33 | 0.91 | 0.24 | 0.60 |
| hsa-miR-6068 | -1.61 | 2.25E-02 | 0.95 | 0.17 | 0.12 | 0.50 |
| hsa-miR-133a-3p | 1.64 | 2.31E-02 | 0.57 | 0.74 | 0.32 | 0.67 |
| hsa-miR-3688-3p | 1.20 | 3.45E-02 | 0.52 | 0.81 | 0.34 | 0.68 |
| hsa-miR-656-3p | 1.34 | 3.66E-02 | 0.67 | 0.65 | 0.32 | 0.64 |
| hsa-miR-659-5p | -1.44 | 3.97E-02 | 0.90 | 0.41 | 0.31 | 0.63 |
| hsa-miR-122b-5p | 1.01 | 4.06E-02 | 0.77 | 0.48 | 0.25 | 0.60 |
| hsa-miR-4473 | -1.42 | 4.07E-02 | 0.87 | 0.46 | 0.33 | 0.63 |
| hsa-miR-33a-5p | 1.31 | 4.61E-02 | 0.52 | 0.81 | 0.34 | 0.65 |
| hsa-miR-194-5p | 0.53 | 9.44E-02 | 0.52 | 0.76 | 0.28 | 0.60 |

